# Towards genuine three-dimensional diffusion imaging with physiological motion compensation

**DOI:** 10.1101/2024.09.08.611927

**Authors:** Yishi Wang, Dehe Weng, Jieying Zhang, Tianyi Qian, Wenzhang Liu, Kun Zhou, Yanglei Wu, Baogui Zhang, Qing Li, Jing Jing, Zhe Zhang

## Abstract

**Purpose:** We aim to implement a 3D DWI sequence and show its usage on patients with new ischemic lesions.

**Materials and Methods:** The proposed 3D DWI sequence was implemented by integrating second-order gradient moment nulling (M2) and cardiac motion synchronization (Sync). All data were acquired on a 3T MAGNETOM Prisma scanner (Siemens Healthcare, Erlangen, Germany) using a 64 channel head and neck coil. 21 healthy volunteers underwent 3D DWI scans at 0.9 mm isotropic resolution using four motion compensation methods for comparison: no compensation (M0), M2 only, Sync only and the proposed M2+Sync method. 2D phase variation maps with different motion compensation methods were also acquired for one subject to illustrate the mechanism of the proposed method. A ghost-to-signal ratio (GSR) and blurring index was defined and compared among the four methods with repeated measures ANOVA and Tukey’s test. 3D DWI was compared with 2D DWI for ADC quantification. Image quality and ischemic lesion conspicuity were evaluated with 12 patients after endovascular treatment.

**>Results:** Whole brain 3D DWI was achieved at 0.9 mm isotropic resolution within 5 minutes using the proposed sequence. M2+Sync achieved the lowest level of GSR and blurring along the slice direction. ADC quantification showed no statistically significant difference between M2+Sync compared to 2D DWI. 3D DWI showed similar image quality, higher lesion conspicuity and counts compared to 2D DWI.

**Conclusion:** Direct 3D DWI can be achieved by the combination of second order gradient moment nulling and cardiac synchronization.

## Introduction

Enhancements in spatial resolution for diffusion MRI (dMRI) of the brain hold the potential to discern minute structures, enabling a finer investigation of neuroanatomical features and detailed architecture. While three-dimensional (3D) acquisition has the advantages of achieving high resolution and signal-to-noise ratio (SNR) and has been widely utilized for various MRI sequences, such as whole brain T1-weighted imaging, T2-weighted imaging, and functional MRI, achieving 3D diffusion weighted imaging (3D DWI) in vivo has been constrained by a few technical limitations. Achieving a true 3D acquisition necessitates exciting the entire volume and acquiring the 3D k-space through multiple acquisitions. While assembling different parts of k-space from various acquisitions is relatively straightforward for simple T1 or T2 weighted imaging sequences, DWI poses a significant challenge due to physiological motion-induced phase variations. These variations can degrade image quality in multi-shot acquisitions if not adequately addressed.

One approach to tackle phase variations is 3D nonlinear phase correction (1), which requires time-consuming 3D phase navigator acquisition. Multi-slab acquisition offers a balance between 3D acquisition and scan efficiency (2,3). However, multi-slab remains a hybrid approach that falls short of achieving true 3D k-space encoding and causes slab boundary artifacts (4–7) and extensive efforts have been invested in mitigating these artifacts (4–6).

Several methods effectively mitigate the effect of physiological motion during data acquisition rather than relying on post-acquisition correction. Cardiac synchronization has demonstrated its effectiveness in reducing motion sensitivity while retaining residual artifacts (8–11). Gradient moment nulling has proven highly effective for reducing signal loss of myocardium in cardiac DWI with 2D single-shot sequences (12–16). In theory it can eliminate the phase accumulation caused by physiological motion to a certain order, such as position (zeroth order), velocity (first order) and acceleration (second order). In this study, we propose a 3D DWI method, which combines cardiac motion synchronization and second-order gradient moment nulling to tackle phase variations caused by physiological motion.

Previous studies have reported that new ischemic brain lesions detected on DWI are common after endovascular procedures such as balloon angioplasty and stent placement (17,18). These lesions are typically small (<3 mm) (18) and understanding the frequency, locations, and numbers of these lesions is crucial for treatment evaluation and prognosis. High-resolution isotropic DWI has demonstrated superior detection than conventional 4 mm-thick DWI of microinfarcts (19). In this study, the proposed 3D DWI was applied to a group of patients with new ischemic lesions following endovascular treatment to demonstrate its potential clinical value.

## Methods

### Sequence design

The sequence diagram was shown in Figure 1a. The diffusion encoding gradients with second-order gradient moment nulling had identical peak amplitude but different durations, adopted from Welsh’s work. The required gradient durations are related by

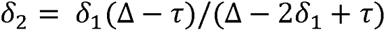

**Figure 1.**
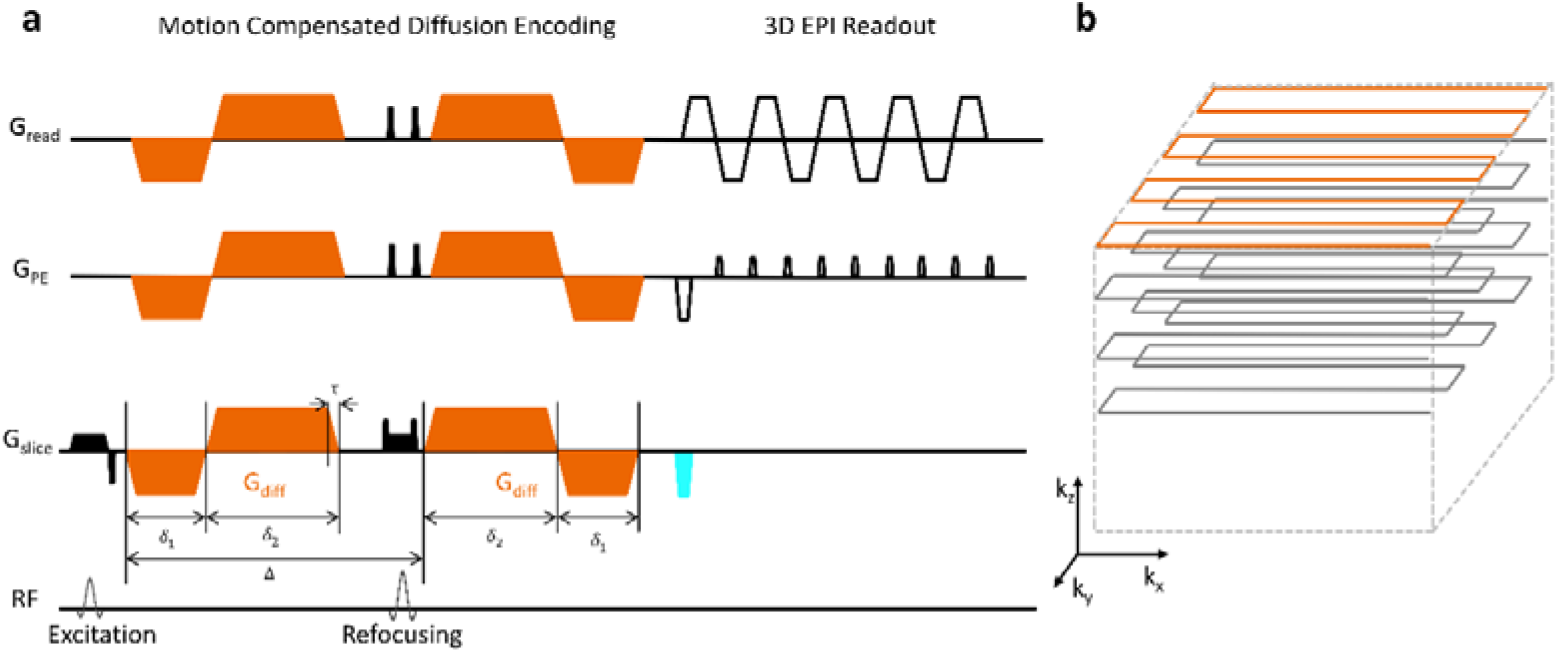
The sequence diagram of 3D DWI using second-order motion compensation and stack of EPI acquisition. **a**, Sequence diagram, the second-order motion compensated diffusion encoding gradients are shown in orange. 2D EPI acquisition is upgraded to 3D acquisition, by adding a slice encoding gradient (cyan trapezoid. 15_1_ and 15_2_ are the durations of the two adjacent gradient lobes in one diffusion encoding block and Δ is the interval between the two blocks. r is the ramping up/down time for each gradient lobe. **b,** 3D stack of EPI trajectory, with one kz plane highlighted in orange, which stands for one portion of k-space covered after each excitation.

**Figure 2.**
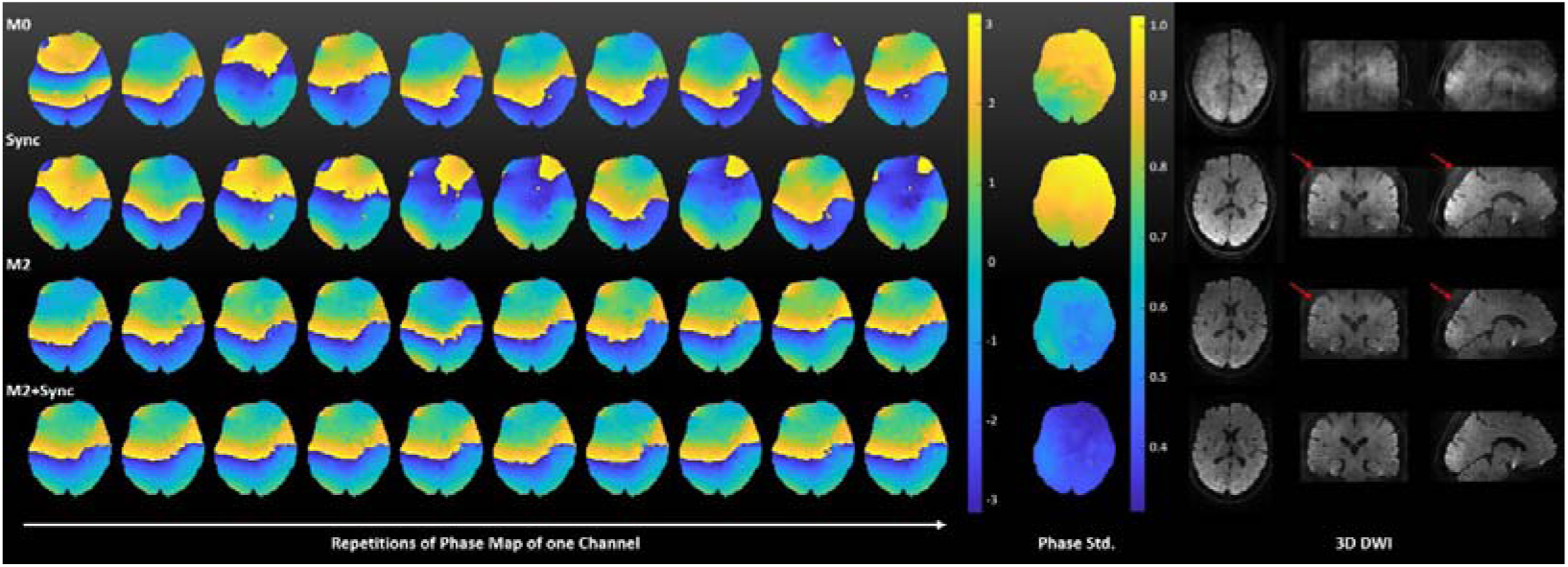
Motion compensation and data consistency. Each row shows the 2D phase maps of 10 repetitions, the standard deviation of the phase maps across 20 repetitions and 3D DWI images. Each row stands for one motion compensation method. In the first row, the data were acquired with without any motion compensation (M0). In the second row, cardiac synchronization was used (Sync). In the third row, second-order gradient moment nulling (M2) was used. In the fourth row, both M2 and Sync were used.

Under these timing conditions, the gradient waveform can generate a diffusion weighting of

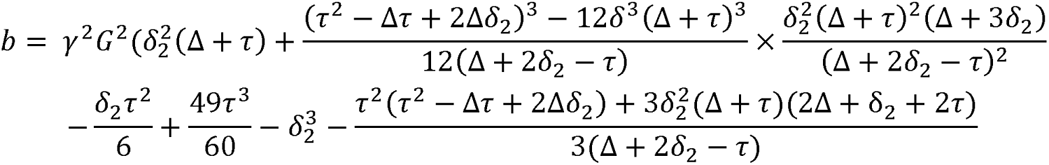

The 3D k-space was acquired with a stack of EPI trajectory shown in Figure 1b. For each TR, after each excitation of the whole 3D volume, only one kz plane is acquired.

### Data acquisition

All data were acquired on a 3T MAGNETOM Prisma (Siemens Healthcare, Erlangen, Germany) using a 64 channel head and neck coil, using foam pads for head fixation. The maximum gradient amplitude was 80 mT/m. 21 healthy volunteers (age 19-38 years, mean age 28.9 years, 7 males) and 12 patients (age 33-75 years, mean age 57.6 years, 3 males) who had ischemic brain lesions after endovascular treatment were recruited for the MRI scan. The inclusion criteria for the patients were: 1, Age between 18 and 80 years old; 2, asymptomatic unruptured cerebral aneurysm treated with stent-assisted coiling/coiling with flow diverter; 3, completion of DWI scan within 48 hours postoperatively. All experiments were performed with IRB guidelines, and written informed consent was obtained from all subjects.

#### Physiological Motion Compensation and Data Consistency

To illustrate the mechanism of the motion compensation strategy, one subject underwent both 2D and 3D DWI scans using four motion compensation methods. For the 2D scans, the four sequences were: 1) monopolar diffusion encoding with 0^th^ moment nulled (M0), 2) M0 with cardiac synchronization using peripheral pulse unit (PPU) trigger (M0 + sync), 3) second order gradient moment nulled (M2) as shown in Figure 1a, and 4) M2 with PPU trigger (M2 +sync). PPU trigger delay was 0ms. Three repetitions of b=0 s/mm^2^ and 20 repetitions of b=1000 s/mm^2^ were performed for each sequence. The diffusion encoding gradient was carried out along the superior-inferior direction, which is most prone to motion induced phase variation for DWI (20–22). Then, the phase maps were calculated using the central region of 64X64 of the k-space. The standard deviation of the phase maps was first calculated across the 20 repetitions for each coil, then averaged across all coils. For the same subject, identical motion compensation methods were used to acquire 3D DWI images.

#### Quantification of motion artifacts

To compare the ghost artifacts level and blurring along the slice direction from the four motion compensation methods, 17 subjects underwent four different sequences at 0.9 mm isotropic spatial resolution.

#### Comparison with 2D Sequence

Nine volunteers underwent a 3D M2+sync and a standard 2D DWI scan for ADC comparison. The patients underwent 2D DWI and 3D M2+sync to compare image quality and lesion conspicuity. All detailed scan parameters were listed in Table 1.

**Table 1.** Scan parameters for DWI sequences.

|  | 2D* | 2D (Clinical) | 3D (Volunteers) | 3D (Patients) |
| --- | --- | --- | --- | --- |
| FOV (mm) | 220×220 | 224×224 | 230×230 | 224×224 |
| Matrix size** | 110×110 | 192×192 | 256×256 | 224×224 |
| Slice thickness(mm) | 4 | 4 | 0.9 | 1.0 |
| Slices | 1 | 27 | 120 | 128 |
| Slice gap (mm) | 1 | 1 | 0 | 0 |
| Bandwidth (Hz/Pixel) | 1818 | 1086 | 888 | 1240 |
| TR (ms or RR interval) | 2000 | 4400 | 750 ms or 1 RR | 1 RR |
| TE (ms) | 63 | 72 | 88 | 72 |
| iPAT | 2 | 2 | 3 | 3 |
| Phase Partial Fourier | 1 | 6/8 | 5/8 | 5/8 |
| Slice Partial Fourier | None | None | 5/8 | 1 |
| Diffusion scheme | Slice direction | 4-Scan Trace | 3-Scan Trace | 4-Scan Trace |
| b value (s/mm<sup>2</sup>) | 0/1000 | 0/1000 | 0/1000 | 0/1000 |
| Scan time (min:sec) | 1:00 | 1:21 | ~4:30 | ~7:30 |

\*For the phase map in data consistency experiment.
\*\*Acquisition matrix = reconstruction matrix

### Image reconstruction

All 3D DWI images were reconstructed with a simple pipeline: after correction of Nyquist ghost (23), GRAPPA (24) was performed to restore the missing data and partial Fourier was reconstructed with a POCS (25) algorithm. The image reconstruction was carried out using MATLAB (The MathWorks, Natick/MA, USA). The 2D images were reconstructed online using the product algorithm.

### ADC Quantification

Seven regions of interest (ROIs) were selected on the ADC maps (Figure 4), including bilateral frontal white matter, corpus collosum, caudate nucleus and thalamus.

### Evaluation of Patient Data

One neurologist (J.J., with >10 years of experience) evaluated the DWI for overall image quality (considering SNR, resolution and motion artifacts) and lesion conspicuity based on a 5-point Likert scale: 1, non-diagnostic; 2, poor but still interpretable; 3, acceptable; 4, good; 5, excellent. The lesion counts were recorded and lesions were considered separate without continuity on the same section and adjacent sections.

## Statistical Analysis

A ghost to signal ratio (GSR) was defined as the mean signal intensity (SI) in the area of the image prone to ghosting relative to SI within the brain along the coronal view. As shown in Figure 3, one ROI was selected as the brain ROI (red circle) and one outside the brain (cyan rectangle) as the ghost ROI. The ROIs were drawn on one slice of the coronal view in the sync image and applied to the same position the other two sequences.

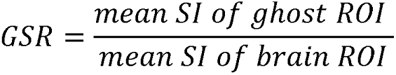

**Figure 3.**
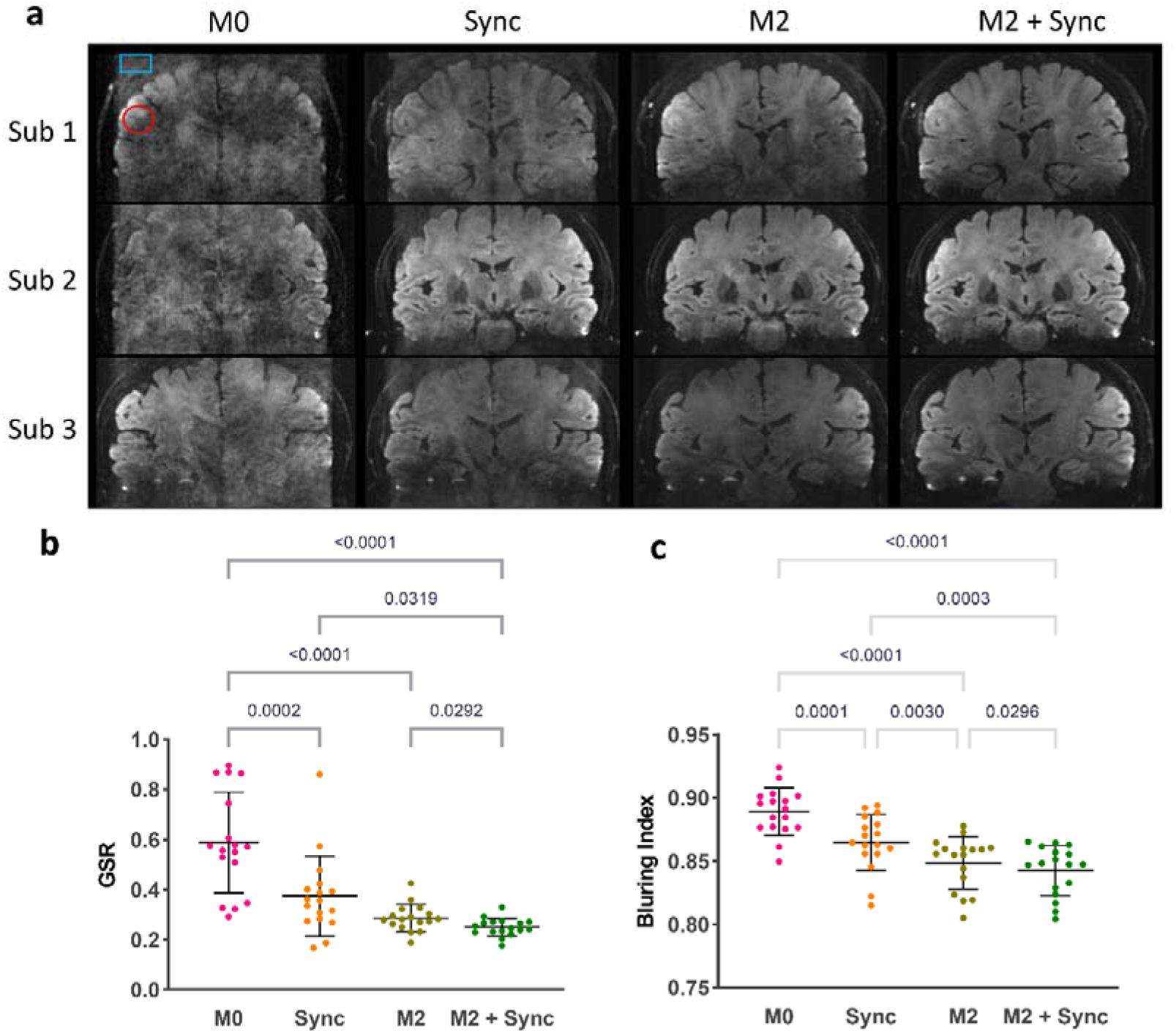
Quantification of motion artifacts. **a**, the coronal view of 3D DWI from three representative subjects with four motion compensation strategies. ROIs were selected to calculate ghost-to-signal ratio. **b,** the GSR of 17 subjects using four motion compensation strategies. c, the blurring index of 17 subjects using four motion compensation strategies.

**Figure 4.**
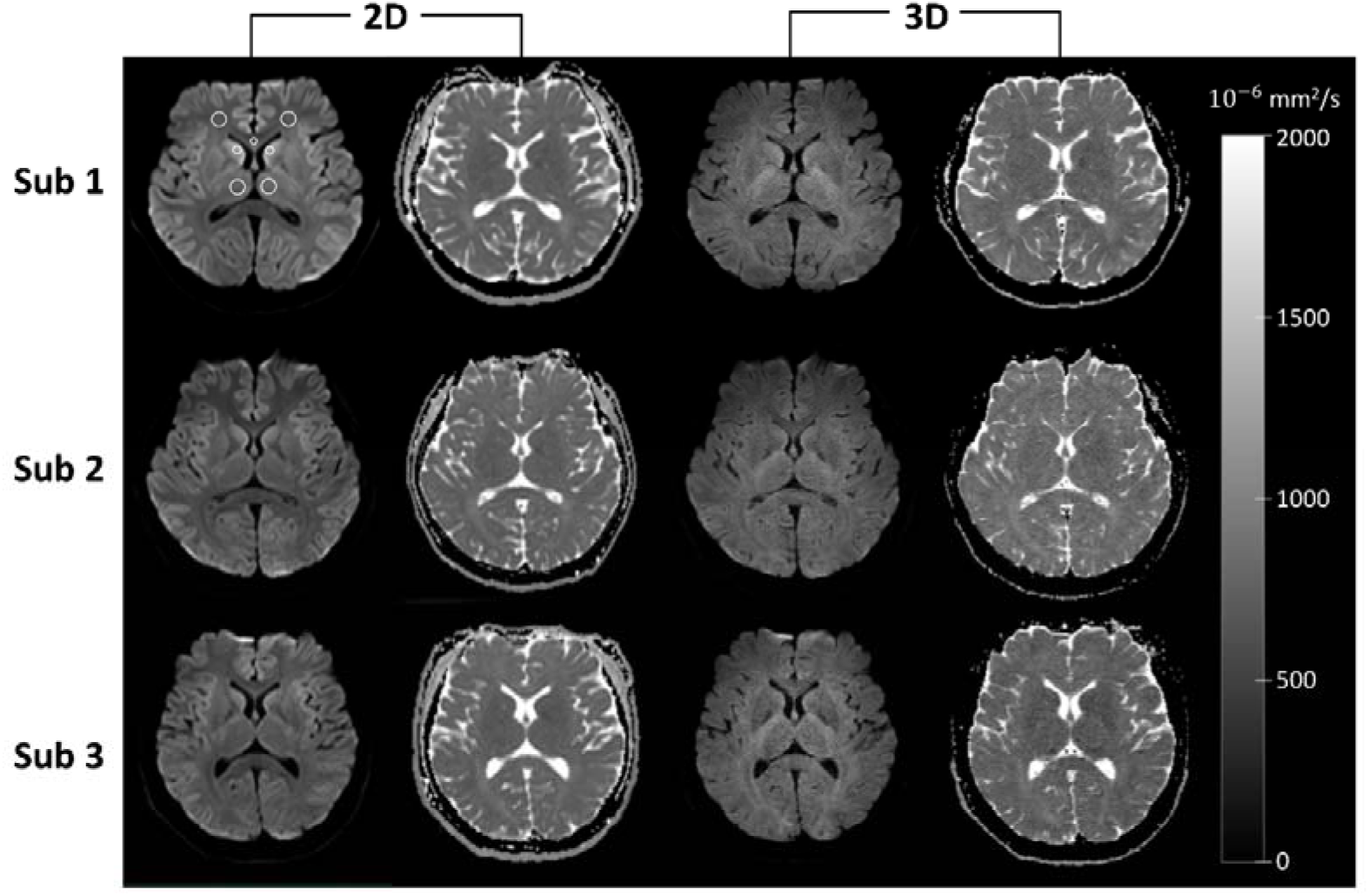
DWI and ADC maps. The DWI and ADC maps of three volunteers were shown for 2D and 3D scans. The white circles are the selected ROIs for ADC quantification.

A blurring index along the z-direction was calculated as correlation coefficients of the image and the image shifted four slices down (26).

First, GSR and blurring index calculated for the 17 subjects passed normality test for all four sequences using Shapiro–Wilk test. Then, repeated measures ANOVA and Tukey’s test was used to compare the GSR and blurring index from different methods. Multiple t-test and Wilcoxon signed rank test was used to compare ADC and lesion counts respectively. All statistical analysis was performed using GraphPad Prism.

## Results

### Physiological Motion Compensation and Data Consistency

Figure 2 shows the phase map of the complex image from 10 out of 20 repeated measurements of the same slice position. The first row shows the data acquired with M0. Notably, the phase patterns exhibited considerable variability across the ten repetitions.

In the second row, only Sync was employed, the phase pattern displayed noticeable differences across repetitions. In the third row, M2 yielded phase patterns with improved uniformity across repetitions. Finally, in the fourth row, M2+Sync resulted in nearly identical phase patterns. The standard deviation of the phase maps, presented on the right side, serves as a quantitative indicator of the different methods’ effects on phase variation. Reduced phase variation signifies a higher degree of data consistency. As illustrated on the right side of Figure 2, 3D DWI images were acquired with identical motion compensation techniques employed in the phase maps. Images acquired without motion compensation exhibited prominent motion artifacts. These artifacts were appreciably mitigated by either Sync or M2, with residual ghost artifacts identified by the red arrows. A combination of the two strategies leads to 3D DWI images with negligible motion artifacts. The structure details are clearly delineated not just in axial, but also reformatted coronal and sagittal views. We can see that, either method alone is effective to improve image quality but not enough, only by a combination of both strategies, can we achieve good 3D DWI.

### Quantification of motion artifacts

Figure 3a shows the coronal view of DWI from three representative subjects, reaffirming the superior image quality achieved through the integrated motion compensation strategy. Figure 3b displays the GSR obtained from all 17 subjects, depicting the variability in GSR values when the Sync strategy was employed individually (0.37 ± 0.15), indicating that its effectiveness was subject-dependent. Conversely, the implementation of either M2 (0.29±0.05) or M2 + Sync (0.25±0.03) yielded significantly reduced GSR values compared to Sync alone. GSR of M0 (0.59±0.20) was the highest. The blurring index for the four sequences were: M0, 0.889±0.019; Sync, 0.865±0.022; M2, 0.849±0.021; M2+Sync, 0.842±0.019.

The statistical analyses for GSR, including Repeated Measures Analysis of Variance (ANOVA) and Tukey’s post hoc multiple comparisons test, demonstrated a statistically significant difference among the four methods (P value for each pair of comparison was shown in Figure 3b), except for Sync versus M2 (P = 0.15), with the combined M2 and Sync strategy exhibiting the lowest level of motion artifacts. For blurring index, there was a statistically significant difference for each pair of comparison (P value for each pair of comparison shown in Figure 3c).

### ADC quantification

Exemplary ADC maps of three subjects were shown in Figure 4. The ADC for each structure was calculated as the mean of the bilateral ROIs and listed in Table 2. There was no statistically significant difference between the ADC of 2D and 3D sequences for the selected structures (P value in Table 2) and they both fell in the range of published ADC values (27–31).

**Table 2.** ADC comparison for 2D DWI and 3D DWI.

| Structure | ADC ($\times 10^{-6}$ mm<sup>2</sup>/s, mean±SD) | | | |
| --- | --- | --- | --- | --- |
|  | 2D | 3D | P value | Literature |
| Frontal WM | 770±25 | 779±50 | >0.99 | 460-900 |
| Caudate Nucleus | 713±14 | 729±55 | >0.99 | 540-930 |
| Corpus Callosum | 772±35 | 832±53 | 0.14 | 600-900 |
| Thalamus | 724±9 | 736±74 | >0.99 | 660-820 |

### Evaluation of Patient Data

Both 2D and 3D DWI reached 4.0 points for the overall image quality. For lesion conspicuity, 2D DWI reached a score of 4.0±0.7 and 3D DWI was 4.9±0.5. The lesion counts were 27.5±35.0 for 2D DWI which was significantly lower than 33.2±37.6 (P<0.0001) for 3D DWI. Figure 5 showed one ischemic lesion (highlighted by the cross) that was clearly seen on the 3D DWI from all three views, but ambiguous on the 2D DWI. Figure 6 showed one lesion that can be identified on both 3D and 2D DWI. However, the shape and outline of the lesion was more well-defined on 3D DWI.

**Figure 5.**
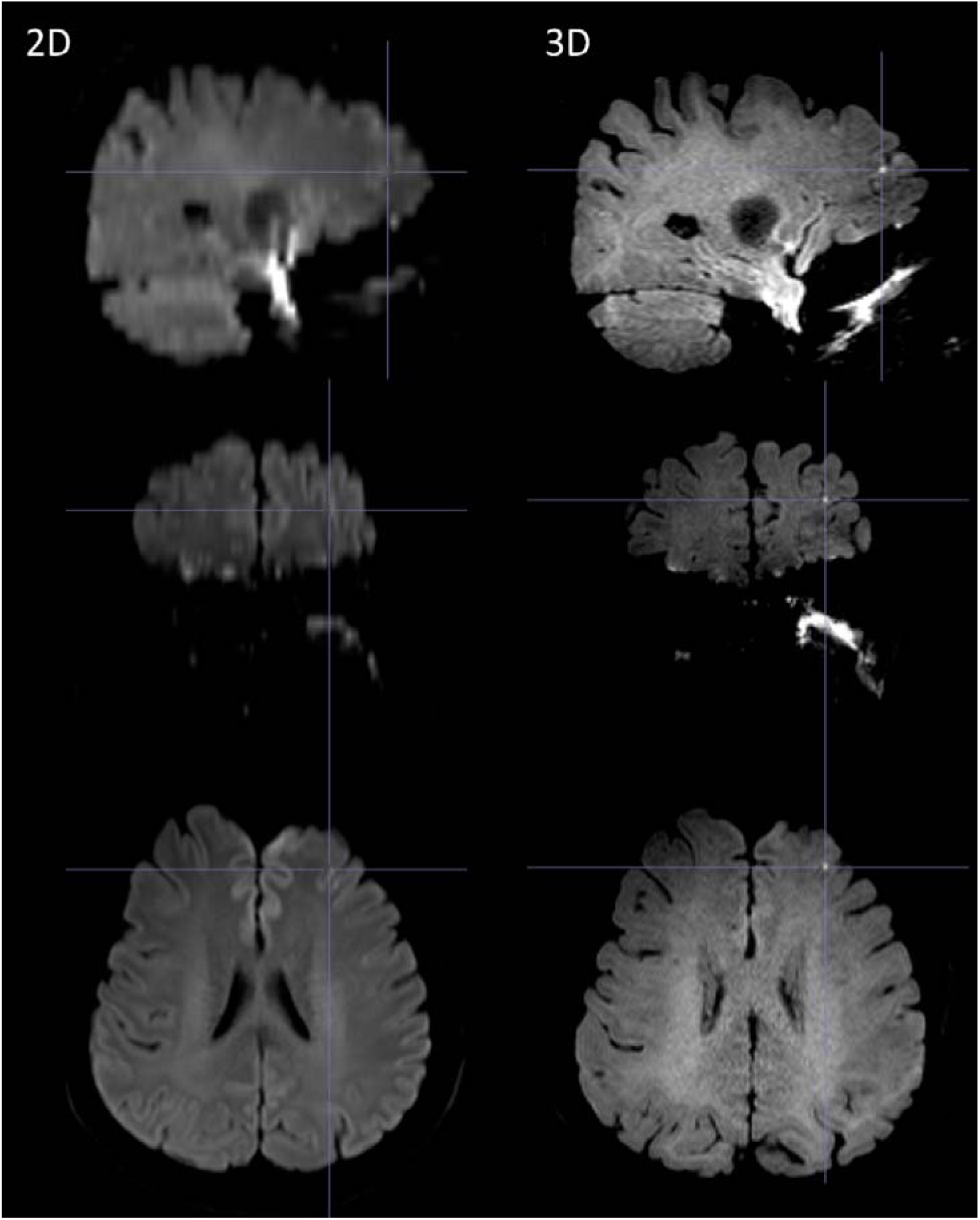
2D and 3D DWI for small ischemic lesion. The cross marked one lesion that was clearly seen on the 3D images but ambiguous on the 2D images.

**Figure 6.**
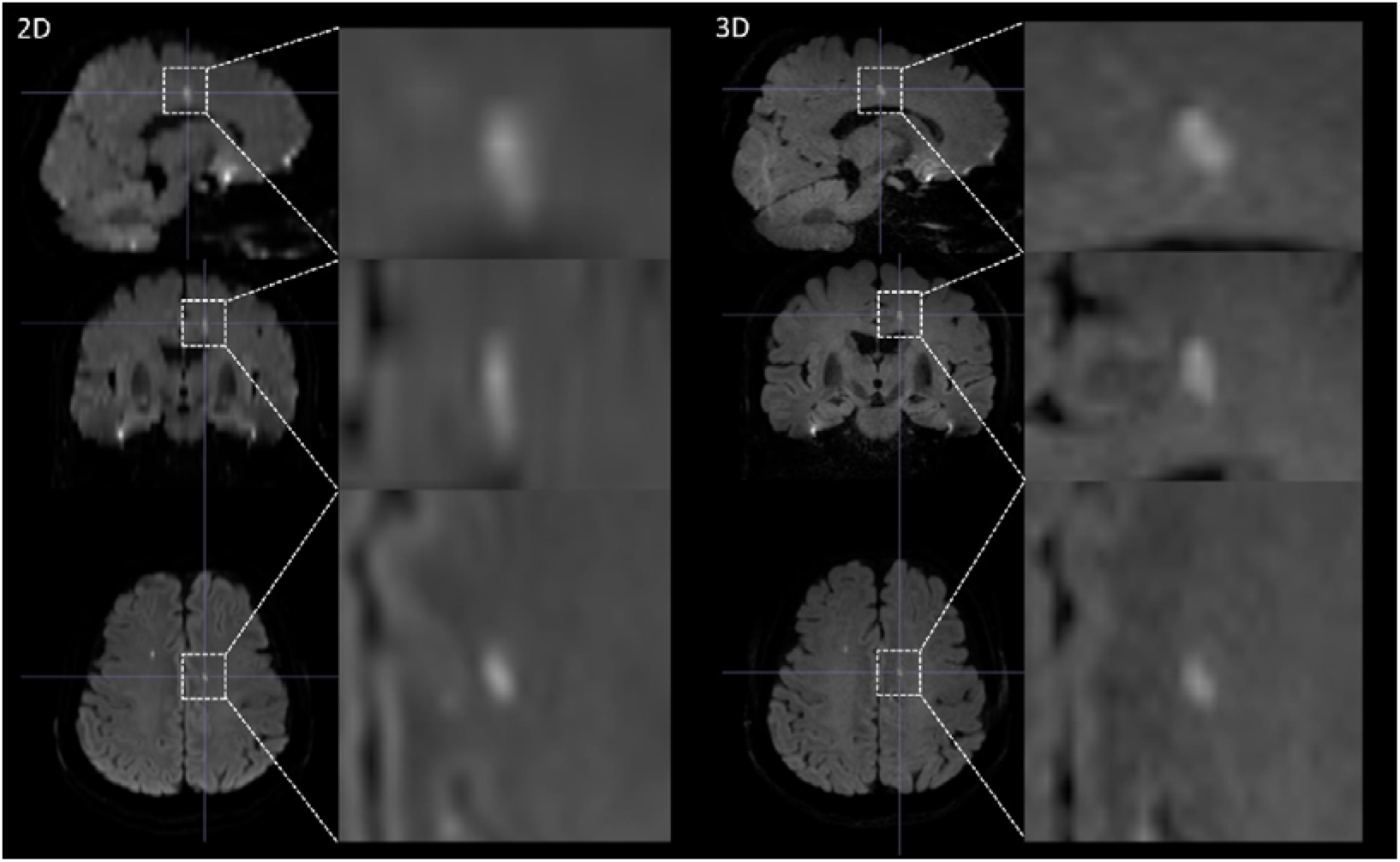
2D and 3D DWI for small ischemic lesion. One lesion that can be identified on both 3D and 2D DWI. The shape and outline of the lesion was more well-defined on 3D DWI as shown in the zoomed in area.

## Discussion and conclusion

In this study, we presented an approach that combines second-order gradient moment nulling and cardiac synchronization to simplify the acquisition and reconstruction of 3D DWI. Moreover, we showcased the ability to obtain sub-millimeter isotropic diffusion imaging using this method. We also evaluated its clinical value with patient data.

A key advantage of this technique is its inherent simplicity both in terms of sequence design, image reconstruction and insensitivity to physiological motion. It obviates the need for phase variation estimation and correction, allowing direct assembly of k-space data from various acquisitions.

For DWI, physiological motion caused by cardiac pulsation, will introduce an extra phase accumulation on the complex valued image. However, the unpredictable motion status during the strong diffusion gradient waveforms leads to a random phase. Second-order gradient moment nulling effectively counteracts phase accumulation due to position, velocity and acceleration, reducing random phase variations. In contrast, cardiac synchronization endeavors near-identical motion conditions for each acquisition, resulting in more consistent phase accumulation. However, the complexity of physiological motion, often exceeding simple linear or cardiac synchronization, underscores the need for their combined usage, consistently minimizing ghost artifacts and blurring along the slice direction across subjects.

In general, 3D MRI provide versatile viewing angles through multi-planar-reconstruction (MPR), offering high-resolution axial and coronal views with a single sequence and minimal image processing. In clinical scan routine, all three views for structural images such as T2 weighted images are typically required for diagnosis. For clinical DWI, axial view is typically used. However, some recent studies proved improved detectability of brain stem ischemia by combining thin slice axial and coronal DWI (32,33). With 3D DWI, high-resolution axial and coronal views can be acquired with only one sequence. Our patient data showed superior conspicuity and fidelity for small ischemic lesions and provided more accurate lesion counts compared to traditional 2D scan. For routine 2D DWI, small ischemic lesions may be missing/blurring due to the partial volume effect from thick slice or low in-plane resolution, or stacked into one slice due to the large slice thickness or even be missed in the slice gap.

For research purposes, techniques like DTI, DSI also necessitate high resolution to overcome partial volume effects. For example, high spatial resolution, rather than angular resolution, is essential for delineation of short association fibers, ranging from 3 to 30 mm, which are crucial for cortical communication (34). High-resolution DTI is also emerging in measuring diffusion anisotropy in the cortex (35–37), which may be useful for probing gray matter micro-structure. Our method is feasible for 3D DTI scan at high spatial resolution (Figures S1, S1). Moreover, although 3D DWI mainly aims for high resolution applications, our technique is straightforward for any moderate to high resolution applications, which maintains scalability as with any other 3D sequences.

Our method has a few limitations. Firstly, M2 leads to a prolonged TE, at the cost of SNR. For example, TE = 88 ms for the high-resolution scan on our scanner using this gradient waveform, while M0 would lead to TE of 20 ms shorter. For a typical clinical 3T scanner with maximum gradient amplitude of 45 mT/m, TE is expected to be about 20 ms longer than the current system (Table S1, Figure S3). However, our work only shows one type of gradient waveform for M2 (14). There are gradient designs which offer time-optimal solutions, with shorter TE for improvement.

Secondly, our method only focused on coping with physiological motion. For bulk motion such as translation and rotation, motion correction technique such as prospective motion correction with fat navigator (38) or optical tracking system (39) is still needed.

Thirdly, using PPU for triggering adds complexity to the scan and make the scan time heart rate dependent. However, this reduces the computing complexity in image reconstruction stage because the shot-to-shot phase variation is mitigated in the data acquisition stage. The short TR (1 RR interval) makes the image more T1 weighted than routine 2D DWI and may reduce the gray matter white matter contrast. This can be mitigated with longer TR (> 1 RR interval). The prolonged scan time can be mitigated with advanced acceleration strategies such as Compressed Sensing (40) to further boost the scan efficiency for 3D DWI.

In conclusion, direct 3D DWI can be achieved by the combination of second order gradient moment nulling and cardiac synchronization. With this technique, submillimeter whole brain isotropic DWI is feasible with about 5 minutes.

## Reference

1. Miller KL, Pauly JM. Nonlinear phase correction for navigated diffusion imaging. Magnetic resonance in medicine 2003;50(2):343–353.

2. Engstrom M, Skare S. Diffusion-weighted 3D multislab echo planar imaging for high signal-to-noise ratio efficiency and isotropic image resolution. Magnetic resonance in medicine 2013;70(6):1507–1514.

3. Frank LR, Jung Y, Inati S, Tyszka JM, Wong EC. High efficiency, low distortion 3D diffusion tensor imaging with variable density spiral fast spin echoes (3D DW VDS RARE). NeuroImage 2010;49(2):1510–1523.

4. Engstrom M, Martensson M, Avventi E, Skare S. On the signal-to-noise ratio efficiency and slab-banding artifacts in three-dimensional multislab diffusion-weighted echo-planar imaging. Magnetic resonance in medicine 2015;73(2):718–725.

5. Van AT, Aksoy M, Holdsworth SJ, Kopeinigg D, Vos SB, Bammer R. Slab profile encoding (PEN) for minimizing slab boundary artifact in three-dimensional diffusion-weighted multislab acquisition. Magnetic resonance in medicine 2015;73(2):605–613.

6. Wu W, Koopmans PJ, Frost R, Miller KL. Reducing slab boundary artifacts in three-dimensional multislab diffusion MRI using nonlinear inversion for slab profile encoding (NPEN). Magnetic resonance in medicine 2016;76(4):1183–1195.

7. Parker DL, Yuan C, Blatter DD. MR angiography by multiple thin slab 3D acquisition. Magnetic resonance in medicine 1991;17(2):434–451.

8. Golay X, Jiang H, van Zijl PC, Mori S. High-resolution isotropic 3D diffusion tensor imaging of the human brain. Magnetic resonance in medicine 2002;47(5):837–843.

9. Van AT, Hernando D, Sutton BP. Motion-induced phase error estimation and correction in 3D diffusion tensor imaging. IEEE Trans Med Imaging 2011;30(11):1933–1940.

10. Jung Y, Samsonov AA, Block WF, Lazar M, Lu A, Liu J, Alexander AL. 3D diffusion tensor MRI with isotropic resolution using a steady-state radial acquisition. Journal of magnetic resonance imaging : JMRI 2009;29(5):1175–1184.

11. Dietrich O, Heiland S, Benner T, Sartor K. Reducing motion artefacts in diffusion-weighted MRI of the brain: efficacy of navigator echo correction and pulse triggering. Neuroradiology 2000;42(2):85–91.

12. Keller PJ, Wehrli FW. Gradient moment nulling through the N11th moment. Application of binomial expansion coefficients to gradient amplitudes. Journal of Magnetic Resonance (1969) 1988;78(1):145-149.

13. Pipe JG, Chenevert TL. A progressive gradient moment nulling design technique. Magnetic resonance in medicine 1991;19(1):175–179.

14. Welsh CL, DiBella EV, Hsu EW. Higher-Order Motion-Compensation for In Vivo Cardiac Diffusion Tensor Imaging in Rats. IEEE Trans Med Imaging 2015;34(9):1843–1853.

15. Das A, Kelly C, Teh I, Stoeck CT, Kozerke S, Chowdhary A, Brown LAE, Saunderson CED, Craven TP, Chew PG, Jex N, Swoboda PP, Levelt E, Greenwood JP, Schneider JE, Plein S, Dall’Armellina E. Acute Microstructural Changes after ST-Segment Elevation Myocardial Infarction Assessed with Diffusion Tensor Imaging. Radiology 2021;299(1):86–96.

16. Stoeck CT, von Deuster C, Genet M, Atkinson D, Kozerke S. Second-order motion-compensated spin echo diffusion tensor imaging of the human heart. Magnetic resonance in medicine 2016;75(4):1669–1676.

17. Hou Z, Jing J, Yan L, Zhang Z, Fu W, Liu J, Yu Y, Jiang L, Yang J, Wang Y, Miao Z, Lou X, Ma N. New Diffusion Abnormalities Following Endovascular Treatment for Intracranial Atherosclerosis. Radiology 2023;307(4):e221499.

18. Cronqvist M, Wirestam R, Ramgren B, Brandt L, Nilsson O, Saveland H, Holtas S, Larsson EM. Diffusion and perfusion MRI in patients with ruptured and unruptured intracranial aneurysms treated by endovascular coiling: complications, procedural results, MR findings and clinical outcome. Neuroradiology 2005;47(11):855–873.

19. Misquitta K, Daou M, Conklin J, Liao C, Setsompop K, Poublanc J, Shirzadi Z, MacIntosh BJ, Tomlinson G, Cohn M, Aviv RI, Silver FL, Mandell DM. Detecting Silent Acute Microinfarcts in Cerebral Small Vessel Disease Using Submillimeter Diffusion-Weighted Magnetic Resonance Imaging: Preliminary Results. Stroke 2022;53(7):e251–e252.

20. Enzmann DR, Pelc NJ. Brain motion: measurement with phase-contrast MR imaging. Radiology 1992;185(3):653–660.

21. Greitz D, Wirestam R, Franck A, Nordell B, Thomsen C, Ståhlberg F. Pulsatile brain movement and associated hydrodynamics studied by magnetic resonance phase imaging. Neuroradiology 1992;34(5):370–380.

22. Poncelet BP, Wedeen VJ, Weisskoff RM, Cohen MS. Brain parenchyma motion: measurement with cine echo-planar MR imaging. Radiology 1992;185(3):645–651.

23. Maier JK, Epstein FH, XZhou X, Ploetz LE, Licato PE; Method for producing an off-center image using an EPI pulse sequence1995.

24. Griswold MA, Jakob PM, Heidemann RM, Nittka M, Jellus V, Wang J, Kiefer B, Haase A. Generalized autocalibrating partially parallel acquisitions (GRAPPA). Magnetic resonance in medicine 2002;47(6):1202–1210.

25. Haacke EM, Lindskogj ED, Lin W. A fast, iterative, partial-fourier technique capable of local phase recovery. Journal of Magnetic Resonance 1991;92(1):126–145.

26. Shou Q, Shao X, Wang DJJ. Super-Resolution Arterial Spin Labeling Using Slice-Dithered Enhanced Resolution and Simultaneous Multi-Slice Acquisition. Front Neurosci 2021;15:737525.

27. Johanna H, Lauri S, Jussi P, Oili S, Aki K, Markku K, Richard ADC, Hannu JA, Turgut T. Diffusion-Weighted MR Imaging in Normal Human Brains in Various Age Groups. American Journal of Neuroradiology 2002;23(2):194.

28. Sener RN. Diffusion MRI: apparent diffusion coefficient (ADC) values in the normal brain and a classification of brain disorders based on ADC values. Computerized medical imaging and graphics : the official journal of the Computerized Medical Imaging Society 2001;25(4):299–326.

29. Klimas A, Drzazga Z, Kluczewska E, Hartel M. Regional ADC measurements during normal brain aging in the clinical range of b values: a DWI study. Clin Imaging 2013;37(4):637–644.

30. Johansson J, Lagerstrand K, Bjorkman-Burtscher IM, Laesser M, Hebelka H, Maier SE. Normal Brain and Brain Tumor ADC: Changes Resulting From Variation of Diffusion Time and/or Echo Time in Pulsed-Gradient Spin Echo Diffusion Imaging. Investigative radiology 2024.

31. Cihangiroglu M, Ulug AM, Firat Z, Bayram A, Kovanlikaya A, Kovanlikaya I. High b-value diffusion-weighted MR imaging of normal brain at 3T. European journal of radiology 2009;69(3):454–458.

32. Baggett M, Helmy D, Chang J, Bobinski M, Assadsangabi R. Added value in stroke imaging: accuracy and utility of additional coronal diffusion-weighted imaging. Clin Radiol 2021;76(10):785 e781-785 e787.

33. Felfeli P, Wenz H, Al-Zghloul M, Groden C, Forster A. Combination of standard axial and thin-section coronal diffusion-weighted imaging facilitates the diagnosis of brainstem infarction. Brain Behav 2017;7(4):e00666.

34. Chen N-k, Guidon A, Chang H-C, Song AW. A robust multi-shot scan strategy for high-resolution diffusion weighted MRI enabled by multiplexed sensitivity-encoding (MUSE). NeuroImage 2013;72:41–47.

35. McNab JA, Polimeni JR, Wang R, Augustinack JC, Fujimoto K, Stevens A, Janssens T, Farivar R, Folkerth RD, Vanduffel W, Wald LL. Surface based analysis of diffusion orientation for identifying architectonic domains in the in vivo human cortex. NeuroImage 2013;69:87–100.

36. Ma Y, Bruce IP, Yeh C-H, Petrella JR, Song AW, Truong T-K. Column-based cortical depth analysis of the diffusion anisotropy and radiality in submillimeter whole-brain diffusion tensor imaging of the human cortical gray matter in vivo. NeuroImage 2023;270:119993.

37. Wu W, Poser BA, Douaud G, Frost R, In MH, Speck O, Koopmans PJ, Miller KL. High-resolution diffusion MRI at 7T using a three-dimensional multi-slab acquisition. Neuroimage 2016;143:1–14.

38. Tisdall MD, Hess AT, Reuter M, Meintjes EM, Fischl B, van der Kouwe AJ. Volumetric navigators for prospective motion correction and selective reacquisition in neuroanatomical MRI. Magnetic resonance in medicine 2012;68(2):389–399.

39. Callaghan MF, Josephs O, Herbst M, Zaitsev M, Todd N, Weiskopf N. An evaluation of prospective motion correction (PMC) for high resolution quantitative MRI. Front Neurosci 2015;9:97.

40. Lustig M, Donoho D, Pauly JM. Sparse MRI: The application of compressed sensing for rapid MR imaging. Magnetic resonance in medicine 2007;58(6):1182–1195.

