## Supplemental Table 1, Figure 1, 2, 3 for "Towards genuine three-dimensional diffusion imaging with physiological motion compensation"

Table S1. Comparison of sequence parameters for different spatial resolutions on two 3T scanners.

| Resolution (mm) | | TE (ms) | | | Bandwidth (Hz/Pixel) | |
| --- | --- | --- | --- | --- | --- | --- |
|  | Prisma 3-scan Trace | | Skyra 4-scan Trace | Prisma 3-scan Trace | | Skyra 4-scan Trace |
| 0.9 | 88 | | 105 | 1220 | | 888 |
| 1.0 | 75 | | 102 | 1208 | | 946 |
| 1.5 | 65 | | 95 | 1934 | | 1218 |

Other identical sequence parameters were: FOV = 230 mm $\times$ 230 mm, maximum b value = 1000 s/mm^2^, iPAT = 3 in PE direction, Phase Partial Fourier = 5/8. The maximum gradient amplitude was 80 mT/m for Prisma and 45 mT/m for Skyra. The maximum gradient slew rate was 200 mT/(m*ms) for Prisma and 180 mT/(m*ms) for Skyra. The bandwidth for each protocol was set to obtain the shortest echo spacing (to reduce distortion) while keeping the bandwidth as small as possible for best SNR.

**3D DTI using M2+Sync**

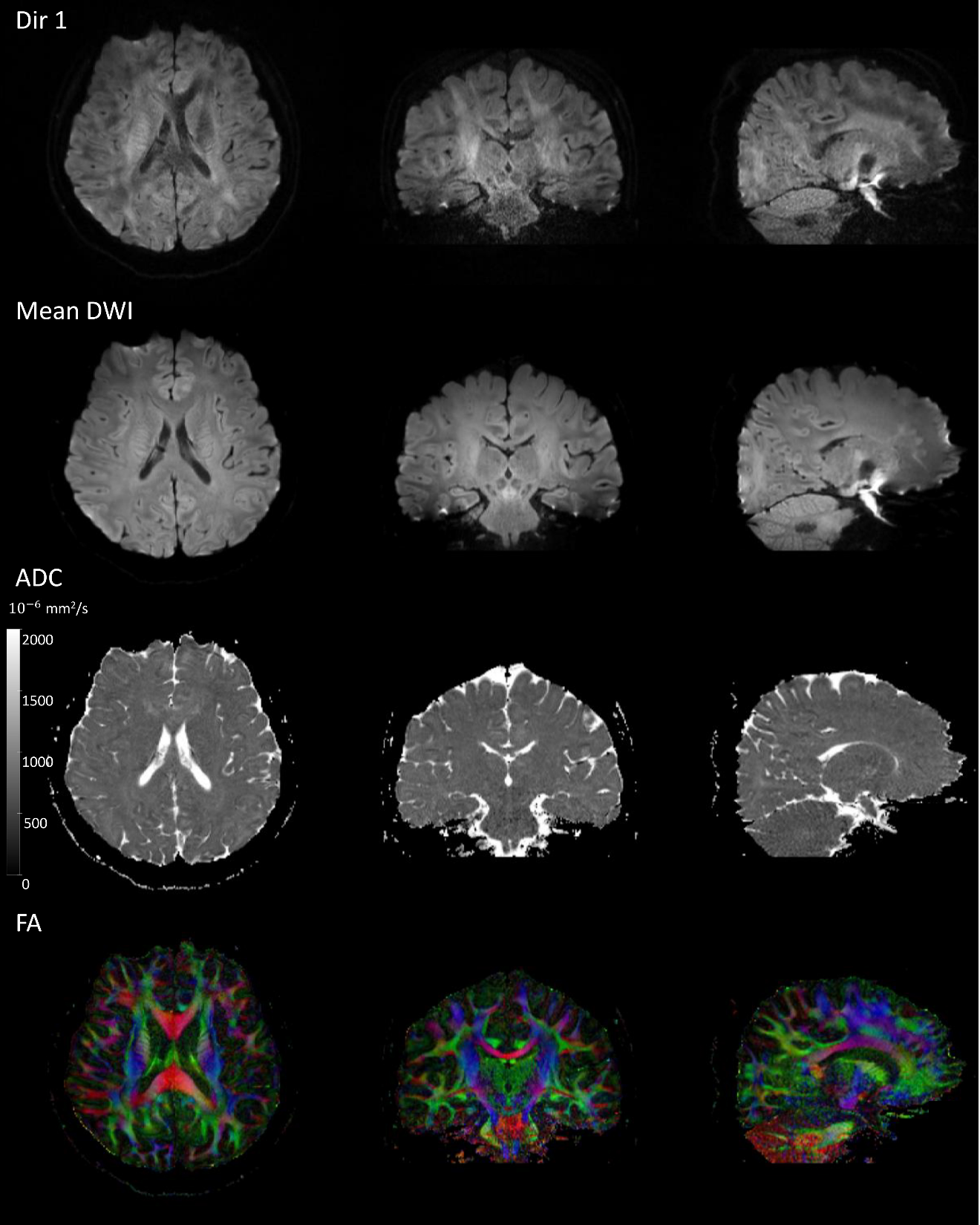

**Figure S1.** 3D DTI of one healthy volunteer (Male, 33 years old) at 0.9 mm isotropic resolution. 1 b=0 s/mm^2^ and 12 directions of DWI using b=800 s/mm^2^. The acquisition time was approximately 16 min.

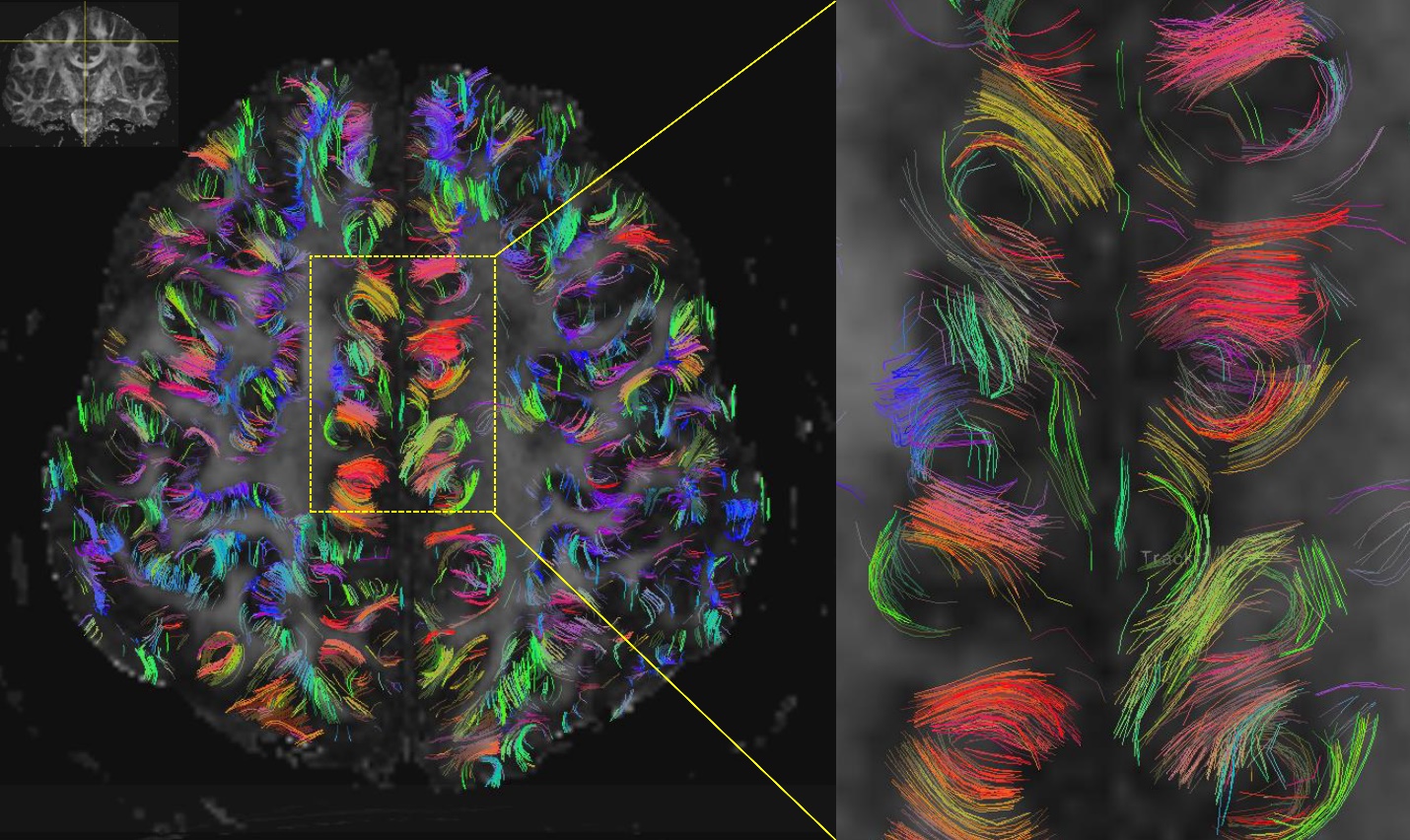

**Figure S2**. Short range fiber tracking using 3D DTI. The same data as in Figure S1, showing highly curved U-fibers.

**3D DWI data on Prisma scanner while mimicking the expected TEs of a Skyra scanner.**

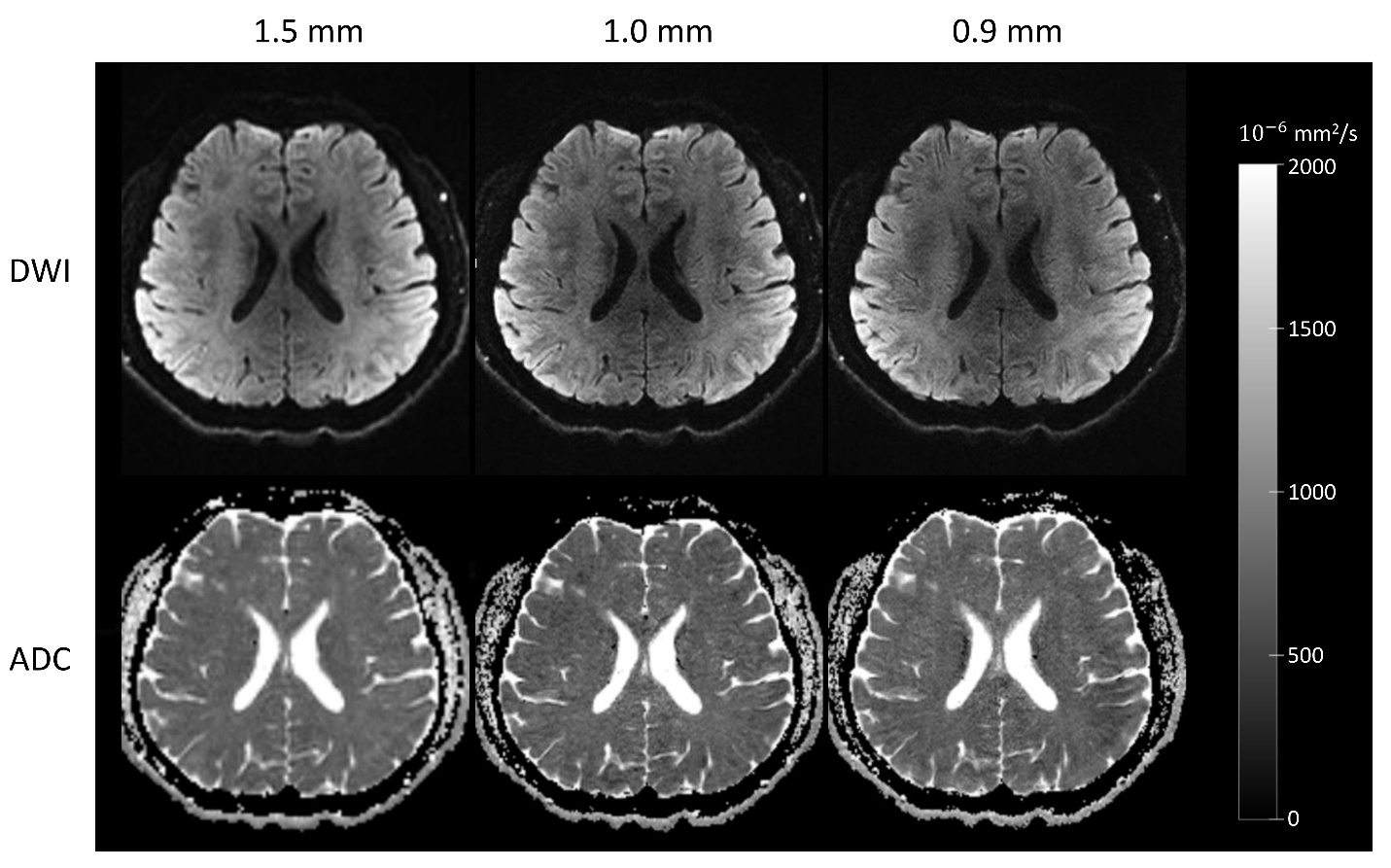

Figure S3. The DWI at b=1000 s/mm^2^ and ADC maps of 3D acquisitions using the expected TEs and bandwidths of a weaker gradient system. The spatial resolution were 1.5 mm, 1.0 mm and 0.9 mm isotropic respectively. The corresponding TEs were 95 ms, 102 ms and 105 ms respectively. The bandwidths were 888 Hz/pixel, 946 Hz/pixel, 1218 Hz/pixel, respectively. 4-scan trace was used for each acquisition in order to obtain the shortest TE.
